# A deep convolutional neural network trained for lightness constancy is susceptible to lightness illusions

**DOI:** 10.1101/2025.11.10.687742

**Authors:** Jaykishan Patel, Alban Flachot, Javier Vazquez-Corral, David H. Brainard, Thomas S. A. Wallis, Marcus A. Brubaker, Richard F. Murray

## Abstract

Human viewers are able to perform tasks that depend on accurate estimates of surface reflectance, even across large changes in illumination and context. This is a remarkable ability, and successful image-computable models of how the visual system achieves this have remained elusive. Recently, deep convolutional neural networks (CNNs) have been developed that are adept at estimating surface reflectance. Here we evaluated one such network as a starting point for a new model of human lightness perception by testing whether it was susceptible to a range of classic lightness illusions. We implemented a CNN and trained it via supervised learning to estimate surface reflectance at each pixel in grayscale, rendered images of geometric objects. We examined the network’s output on several illusions, including the argyle, Koffka, snake, simultaneous contrast, White’s, and checkerboard illusions, as well as control figures. We included variants where low-luminance regions important to the illusions were generated either by low reflectance or by cast shadows. For comparison, we carried out a lightness matching experiment with human observers using the same stimuli, and also examined the outputs of three classic lightness and brightness models. The CNN largely removed lighting effects such as shading and shadows, and produced good reflectance estimates on a test set. It also qualitatively predicted the illusions perceived by humans in most cases, the exceptions being White’s and checkerboard illusions. The CNN outperformed classical models, both at estimating reflectance and at tracking human lightness matches. These findings support a normative view of lightness perception and highlight the promise of deep learning models in this area.

## Introduction

The human visual system is able to use 2D retinal images to make accurate judgements of features of the external world, even in complex and dynamic environments. In particular, a topic of long-standing interest is how the brain creates representations of achromatic surface color (Adelson, 2000; Gilchrist, 2006; Murray, 2021).

One common measure of achromatic surface color is *reflectance*, the proportion of spectral power in the visible wavelength range that is reflected by a surface (McCluney, 1994). Reflectance is a physical property. *Lightness* is the perceptual correlate of reflectance, and lightness constancy is the ability to maintain a stable percept that is correlated with reflectance despite varying lighting conditions and context (Gilchrist, 2006). Having a perceptual correlate of reflectance is useful for many tasks, such as object recognition, scene interpretation, and identifying materials. We perceive lightness at a glance, but this ability has proven difficult to understand and model. Many combinations of shape, lighting, and material properties can produce any given retinal image, so creating a representation that is strongly correlated with surface reflectance is a challenging computational problem (Belhumeur, Kriegman, & Yuille, 1999). Decades of perceptual experiments have revealed some of the image features that guide lightness perception, but a broadly successful, predictive, image-computable model of lightness has remained elusive (Adelson, 2000; Gilchrist, 2006; Murray, 2021).

Inferring reflectance and illumination from an image is also a classic problem in computer vision, where it is called ‘intrinsic image decomposition’ (Barrow, Tenenbaum, Hanson, & Riseman, 1978). Here too, substantial progress has been made over many years of research, but until recently advances were mostly made in limited stimulus domains, and inferring surface reflectance in arbitrary, complex scenes has been a challenge (Barron & Malik, 2015; Rother, Kiefel, Zhang, Schölkopf, & Gehler, 2011; Shen, Yeo, & Hua, 2013; Weiss, 2001).

In recent years, deep learning methods have been applied to intrinsic image decomposition, with promising results (Li & Chen, 2015; Li et al., 2021; Sato et al., 2024; Y. Yu & Smith, 2018). These studies have used many kinds of training data, from renderings of simple objects to web-crawled images of real scenes, and have achieved impressive performance at recovering surface reflectance in novel images. In some cases they have been successful enough to support downstream tasks such as object insertion (Li et al., 2021; B. Yu et al., 2023).

These results raise interesting questions about deep learning approaches to lightness perception and intrinsic image decomposition. Are these networks plausible starting points for models of human lightness perception? What image features and computations do they use to infer surface reflectance? Earlier work has suggested that seemingly idiosyncratic properties of human lightness perception are in fact adaptations to the statistics of natural images (Murray, 2013, 2020; Purves & Lotto, 2010); do these networks, which learn from natural images, show these idiosyncratic behaviours as well? In particular, do these networks reproduce the strong lightness illusions that have been discovered in work on human vision?

Here we begin to address these questions, by testing whether a convolutional neural network (CNN) that has been trained to infer surface reflectance is subject to many of the same lightness illusions as human observers. We trained the network to infer reflectance from renderings of scenes of randomly placed and textured geometric objects. We chose this simple training set instead of real or photo-realistic images because we wished to test the hypothesis that important features of lightness perception can emerge from generic features of 3D scenes, such as shadows, shading, and occlusion, rather than more detailed properties of real scenes such as specific materials or common shapes. We compared network performance to human performance, by evaluating the network’s lightness matching behaviour under the computational linking hypothesis that two stimuli are matched for the network when their inferred reflectances match. We performed this analysis for several well-known lightness illusions, and collected new human lightness matching data for the same set of illusions. We also examined how well three classic lightness and brightness models predicted human performance for these illusions.

We had two main goals with this work. The first was theoretical. We wished to evaluate the idea that many robust features of human lightness perception are byproducts of the visual system’s attempt to produce stable perceptual correlates of surface reflectance from ambiguous retinal images. In particular, we wished to test the idea that many lightness illusions are not simply failures of perception, but can emerge as byproducts of an adaptive, data-driven strategy that recovers surface reflectance from retinal images of complex 3D scenes. We tested whether a model optimized for reflectance estimation in naturalistic settings exhibits illusion-like behavior as a consequence of its learning, thereby providing a computational perspective on why such illusions may arise in human vision. Previous studies have tested models that incorporate the ambiguity of retinal images, but they have been limited to simple 2D images (Allred & Brainard, 2013; Murray, 2020). Our goal was to exploit the power of deep learning methods to extend this approach.

Our second goal was practical. We wished to test whether a specific architecture of convolutional neural network could be trained to estimate reflectance from rendered images, and we also wished to evaluate whether this network could serve as a starting point for a new model of human lightness perception. Experimental work has revealed important properties of lightness perception, but has not resulted in image-computable models that capture a wide range of lightness phenomena. Here we evaluate a type of model that, should it appear promising, can be further developed using newly emerging methods.

For this investigation, we used a network that is a subset of InverseRenderNet, an architecture that has been developed in the intrinsic image decomposition literature (Yu & Smith, 2018). InverseRenderNet is a convolutional U-Net with skip connections, that maps RGB input images to pixelwise estimates of RGB reflectance and illuminance. Our network, which we call IRNet, is a modification of InverseRenderNet that maps achromatic luminance inputs to achromatic albedo outputs. Thus we simply reduced number of input and output channels from three (RGB) to one (luminance input, albedo output), and discarded the decoder that estimates illuminance. We did not use the pre-trained weights for InverseRenderNet, and instead trained IRNet from scratch (details below). Our goal was to examine a representative CNN that has been shown to be successful at estimating surface reflectance, and to compare its performance in a psychophysical task to that of human observers. To encourage further exploration of this modeling framework, we have made code for training and evaluation of the network publicly available (https://github.com/JaykishanPatel/irnet_supporting).

### Previous work

Previous studies have taken many approaches to modelling human judgments of lightness, colour, and brightness. For example, Brainard et al. (2006; see also Brainard and Freeman (1997)) developed Bayesian methods for illuminant estimation and linked these to human judgments of color appearance for a range of visual stimuli that were computer renderings of illuminated scenes. They showed that this general approach could account for both successes and failures of human color constancy. Their work considered only a single geometric arrangement of scenes and assumed that there was only a single illuminant common to all objects within each scene. Allred and Brainard (2013) took a step towards generalizing these ideas to spatially varying illumination in the achromatic case, but did not consider 3D scene structure.

Many other studies have taken quantitative and computational approaches to modelling lightness, colour, and brightness. For reviews see Adelson (2000), Gilchrist (2006), Kingdom (2008, 2011), Brainard and Maloney (2011), Schirillo (2013), and Murray (2021). In the rest of this brief overview, we focus on deep learning models.

Corney and Lotto (2007) trained an artificial neural network to predict reflectance from luminance images. The training and test sets consisted of images from a synthetic ‘dead leaves’ dataset, where each image depicted many overlapping patches under smoothly varying illumination. The trained network accounted for several well-known brightness illusions, and furthermore predicted parametric variations in illusion strength that were consistent with human perception. The network was tiny by current standards (just four hidden-layer neurons), and the training set consisted of simple 2D images, but this early work demonstrated how machine learning methods can be used to examine the relationship between natural scene statistics and visual perception.

Gomez-Villa, Martín, Vazquez-Corral, Bertalmío, and Malo (2020) evaluated several network architectures that were trained for noise removal, deblurring, or both. Network depths ranged from 2 to 21 hidden layers. They found that shallow networks often produced brightness and color illusions, but interestingly, deeper networks did not. Furthermore, the direction of the illusions generated by shallow networks depended on the task they were evaluated on after training: in an estimation task, the illusion predicted by networks was typically consistent with the illusion perceived by human observers, but in a matching task, it was often in the opposite direction. This study provided a useful complication to claims about the relationship between deep networks and human vision, showing how results from networks can depend on details of architecture and task. The authors also showed that these complex, trained networks could be approximated by a simple linear receptive field. In related work, Gomez-Villa, Martín, Vazquez-Corral, Bertalmío, and Malo (2022) showed that a generative adversarial network could create novel stimuli in which human observers perceived lightness illusions.

Kubota, Hiyama, and Inami (2021) examined convolutional networks with one hidden layer, trained for denoising, deblurring, or blurring. They compared the outputs of these networks to results from two previous psychophysical studies of the Munker-White illusion. They found mixed results, with the best matches to psychophysical data generally produced by the network trained to blur images. This is in contrast with other studies reviewed in this section, where denoising and deblurring often produced human-like results.

Mukherjee, Paul, and Ghosh (2024) examined additional network architectures trained for denoising and deblurring, as well as classical image processing methods for these tasks. Like Gomez-Villa et al. (2020), they found that several of these networks predicted well-known brightness illusions. Significantly, they also found that the classical image processing methods did not produce these illusions, which raises questions about what aspects of denoising and deblurring, learned by the networks but not present in classical methods, give rise to the illusions.

In summary, many recent studies have shown that deep learning models trained on low-level image transformations can exhibit lightness or brightness illusions. The present work builds on this literature by developing another image-computable model that is susceptible to lightness illusions, but that differs in key respects. First, the model we present is explicitly trained to estimate surface reflectance from images, rather than to perform image restoration tasks such as denoising or deblurring. This choice reflects the view that lightness perception is shaped by ecological demands, where the visual system uses natural image statistics to estimate perceptual correlates of latent physical properties, such as reflectance, from ambiguous luminance inputs. By training a model to solve this inverse problem directly, we aim to test whether this objective alone can produce human-like perceptual behaviors, including susceptibility to lightness illusions. Second, we compare the model’s output directly to human psychophysical data collected using the same rendered stimuli, allowing a direct, stimulus-matched comparison. This type of comparison has rarely been performed in the literature.

## Methods

### Experiment with human observers

#### Participants

Eleven naive observers participated in the experiment (five male, six female; ages 20–30 years). All observers reported normal or corrected-to-normal acuity, gave written informed consent, and were paid for their participation. All procedures were approved by the Office of Research Ethics at York University.

#### Stimuli

We recreated several classic lightness illusions using MATLAB (Figure 1) (The MathWorks Inc., 2022). The illusions were the argyle illusion and its long-range and control variants (Adelson, 1993), the Koffka illusion and its control figure (Koffka, 1935), the Koffka-Adelson illusion (Adelson, 2000), a classical simultaneous contrast illusion (Hess & Pretori, 1894/1970), an articulated simultaneous contrast illusion (Katz, 1935), White’s illusion (White, 1979), the checkerboard assimilation illusion (DeValois & DeValois, 1988), and the snake illusion and its control figure (Adelson, 2000). In the control figures that were included for some illusions, low-level factors such as local luminances surrounding the test patches were largely preserved, but features such as X-junctions that gave evidence of lighting boundaries were disrupted. As a result, models based on local contrast would not predict substantially different illusions in the illusion and control figures.

**Figure 1.**
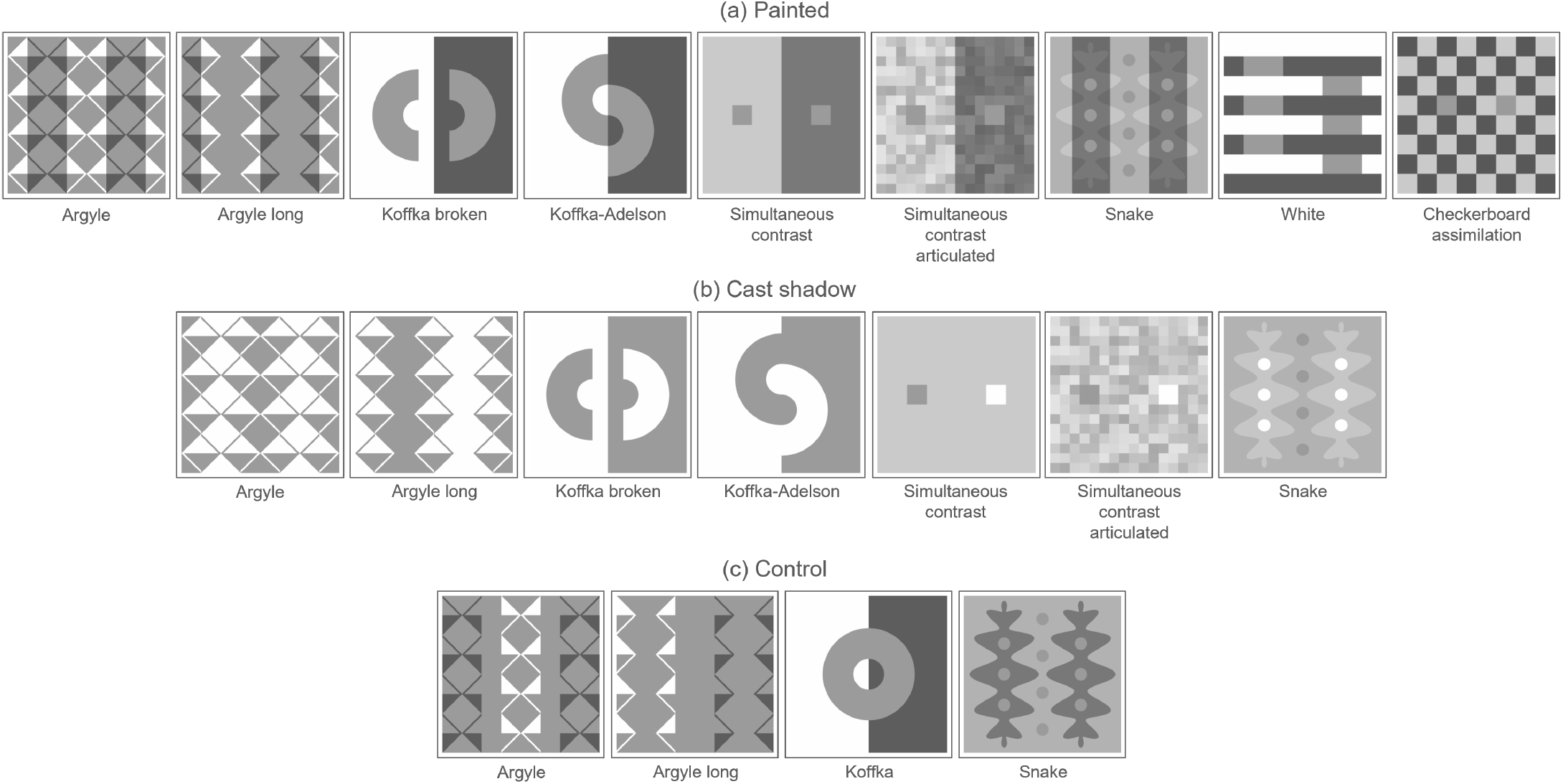
Classic lightness illusions. (a) Painted variants, where dark regions are included in the images shown here. (b) Cast-shadow variants, where dark regions were created by cast shadows in a 3D scene, and are not included in the images shown here. (c) Control variants, where local contrasts are largely preserved but some cues to cast shadows are eliminated.

In each illusion, there were two or more regions that had the same physical luminance, but people typically perceive some of these regions to be lighter than others. Figure 2 shows the equiluminant ‘reference’ and ‘match’ regions (in red and green, respectively) in each image, that we manipulated in the experiment reported below.

**Figure 2.**
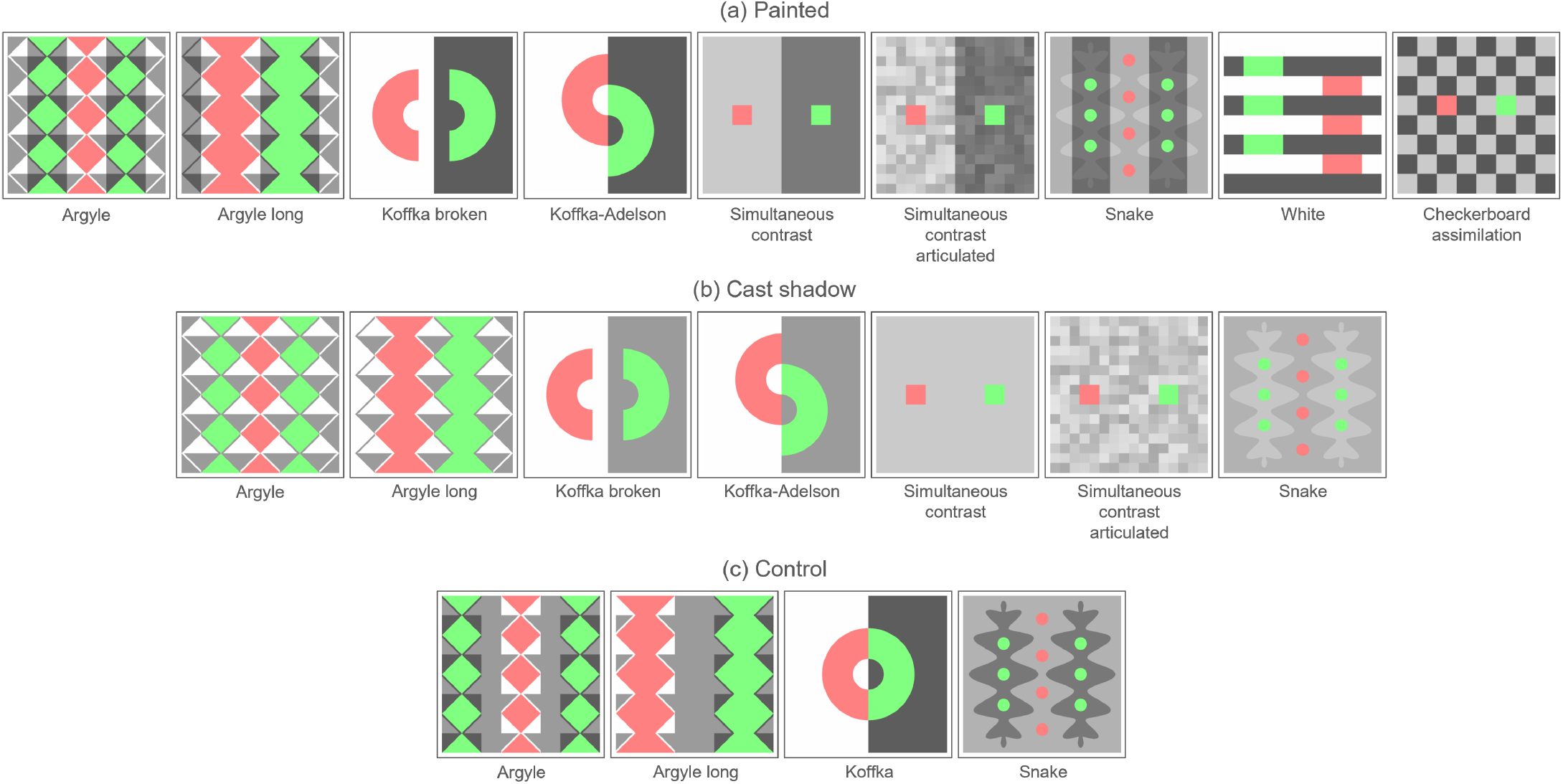
Equiluminant reference and match regions, shown in red and green, respectively, for each image in Figure 1.

In order to make the stimuli similar to the training set for the deep learning network (see below), we rendered these illusions on one side of a cube that was surrounded by several textured objects (Figure 3). We rendered the stimuli using the EEVEE render engine in Blender 2.92 (Blender 2.92 Documentation Team, 2021), at a resolution of 512 *×* 512 pixels.

**Figure 3.**
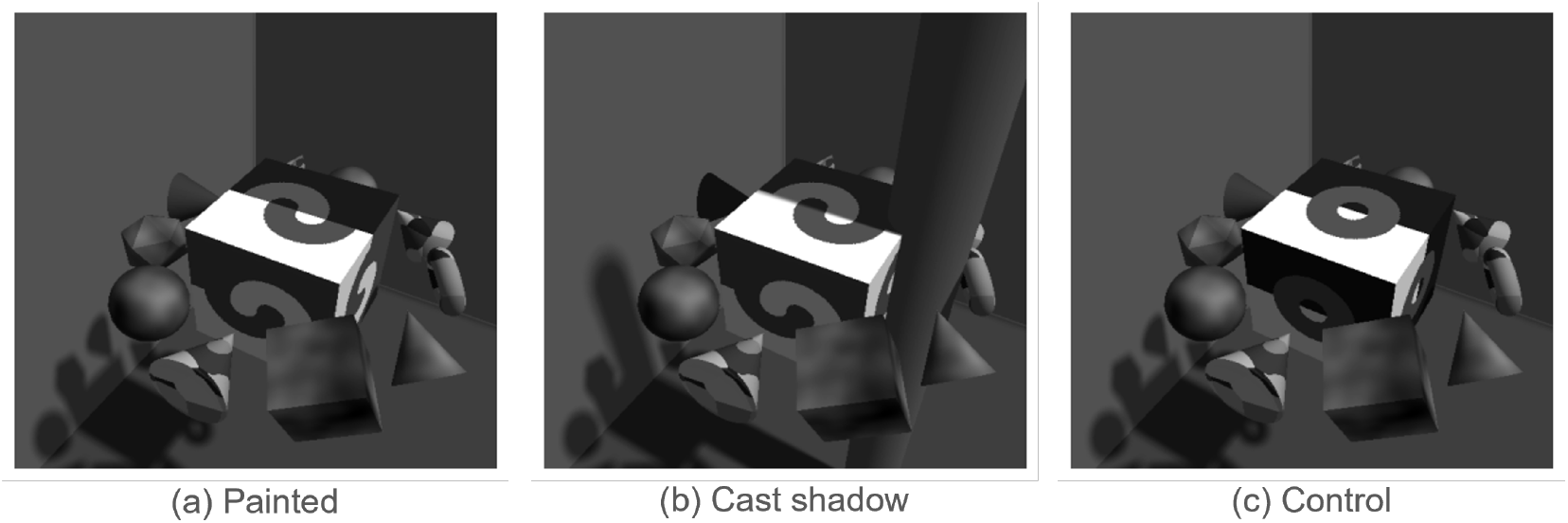
Examples of stimuli in the experiment, showing variants of the Koffka-Adelson illusion on one side of a cube.

In the argyle, Koffka, snake, and simultaneous contrast figures, people typically perceive a test patch in a dark region of the image to be lighter than an equiluminant test patch in a bright region. Perceived shadows may play a role in this phenomenon (Adelson, 1993, 2000; Gilchrist, 2006), so we included two versions of these figures (Figure 3). In the ‘painted’ variant, the classic illusion with light and dark regions was simply rendered on one side of a cube (Figures 1(a), 3(a)). In the ‘cast-shadow’ variant, a figure without the dark regions was rendered on one side of a cube, and the dark regions were created by shadows cast by one or more cylinders in the virtual scene (Figures 1(b), 3(b)). As noted above, for some illusions we also included a control variant, where local contrasts were largely preserved but cues to shadows were disrupted (Figures 1(c), 3(c)). We configured the simulated lighting so that the luminance was the same at each pixel in the painted and cast-shadow variants, except that (a) the cast-shadow variants had a small but visible penumbra along shadow boundaries, whereas the painted variants did not, and (b) in the cast-shadow variants, the scene contained one or more visible cylinders that provided additional evidence that the dark regions were created by cast shadows. The lighting conditions and camera position were the same for all stimuli, except for very small lighting adjustments made in some conditions in order to equate the luminance of the test regions across stimuli (Figure 2), and were within the range used in the training set for the deep learning model (see below).

Stimuli were shown on a 27” iMac (late-2012 model; resolution 1920 *×* 1080 pixels; macOS 10.15), using Python 3.8 and PsychoPy 2024.1.3 (Peirce et al., 2019). We used a Konica Minolta LS-110 photometer and custom software to gamma-correct the monitor. Stimuli measured 12 *×* 12 cm and subtended 6.9 *×* 6.9 degrees of visual angle at the viewing distance of 1.00 m. Overhead lighting was kept on, to reduce the visual impression of light-emitting stimuli in a dark room (Patel, Munasinghe, & Murray, 2018). The displayed luminances ranged from 0.59 to 74 cd/m^2^. Luminance measurements for gamma correction were made with overhead lighting on, under the same viewing conditions used in the experiment, so that any ambient light reflected from the monitor was taken into account during luminance characterization.

#### Procedure

The observer sat at a table that supported the monitor, and the centre of the monitor was approximately at eye level. Head position was stabilized by a chinrest. On each trial, the observer viewed a randomly chosen stimulus depicting a lightness illusion or its control figure (Figure 1) on one side of a cube in a complex scene (Figure 3). The stimulus was shown at the centre of the screen, surrounded by a uniform grey background. At the start of the trial, the reference and match regions (Figure 2) were shown in red and green, respectively, for 1.0 s, to indicate to the observer which regions to match. After this cue, the luminance of the reference region was set to its rendered luminance, which was approximately the same in all images (24.2 *±* 0.1 cd/m^2^), and the luminance of the match region was set to a random luminance within the monitor’s display range. The observer moved a mouse vertically to smoothly adjust the luminance of the match region so that it appeared to have the same reflectance as the reference region. The observer pressed the space bar to indicate that they had completed the match, and then the next trial began. At any time, the observer could click the mouse button to make the reference and match regions briefly flash red and green again, as a reminder of which regions to match. There were 20 stimuli, each repeated 5 times, for a total of 100 trials. The median completion time for the experiment was 20 minutes.

#### Analysis

In each illusion, we chose the reference and match regions such that, from the previous literature, we would expect observers to choose a match luminance that is lower than the reference luminance. Accordingly, we defined the illusion strength to be the reference luminance minus the match luminance. We measured the mean illusion strength for each observer with each illusion.

We excluded one participant’s data from the analysis because their luminance matches were highly variable. The standard deviation of their luminance matches had a mean value of 15.7 cd/m^2^ across conditions, compared to 2.0 cd/m^2^ for the next most variable participant. The mean target patch luminance was 24.2 cd/m^2^, meaning that the standard deviation of this participant’s matches was 65% of the target luminance. Based on this metric, we judged the participant’s data to be unreliable and excluded them from further analysis.

### Experiment with a convolutional network

#### Network architecture

We used Python 3.10 and PyTorch 2.3.1 to implement a subset of the InverseRenderNet network (Paszke et al., 2019; Y. Yu & Smith, 2018). The network was a 30-layer convolutional neural network (CNN) in a U-Net hourglass architecture with skip connections (Figure 4). It had a total of 1.7 *×* 10^7^ weights. The input image was progressively downsampled to a bottleneck layer with 512 channels, and then progressively expanded back up to its original size. The network mapped a 512 *×* 512 luminance image to an equally sized output image that gave pixelwise estimates of achromatic reflectance. We call this architecture IRNet, to indicate that it was derived from InverseRenderNet, but also structurally different in that it had a reflectance readout but no surface orientation readout.

**Figure 4.**
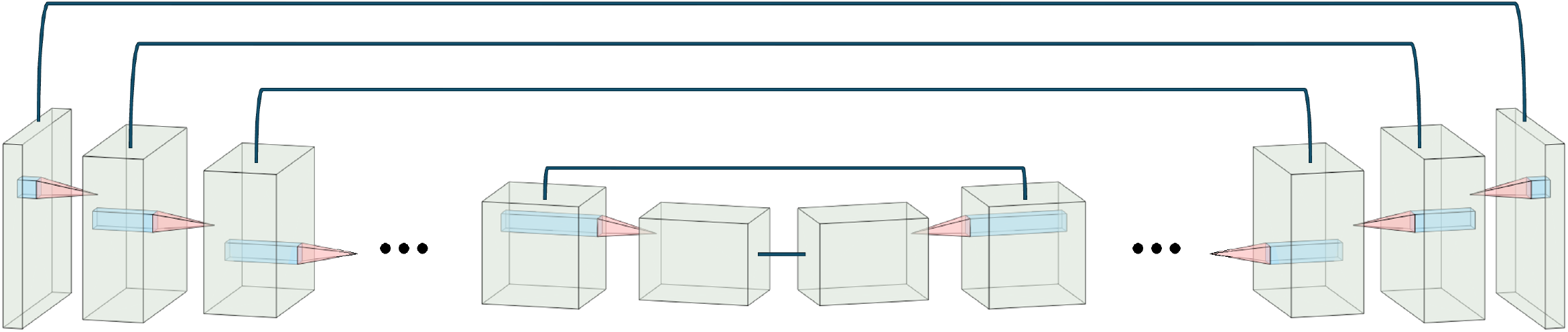
Architecture of IRNet, which is a convolutional U-Net with skip connections. Ellipses indicate additional layers.

#### Training

We rendered 100,000 training image pairs, 5,000 validation pairs, and 5,000 test pairs, using Blender 2.92 with the EEVEE render engine. Each pair consisted of a 512 *×* 512 pixel greyscale image depicting 20 randomly placed geometric objects, and an equally sized image of the reflectance ground truth (Figure 5, columns (a) and (b)). Each object was randomly chosen to be a cube, cylinder, sphere, torus, icosahedron, or cone. Object position, size, and orientation were randomized as well. Each object had a greyscale Lambertian surface reflectance pattern, randomly chosen to be (a) uniform grey with a randomly chosen reflectance in the range [0.05, 0.8], which is approximately the range of reflectances found in natural scenes (Gilchrist, 2006), (b) a randomly generated Voronoi texture, or (c) a randomly generated low-frequency noise texture. Lighting consisted of a directional source and an ambient source, with randomized lighting direction, and randomized directional and ambient intensities. The virtual camera position was also randomized. Details of the randomization of scene parameters can be found in the Supporting Information (link provided below).

**Figure 5.**
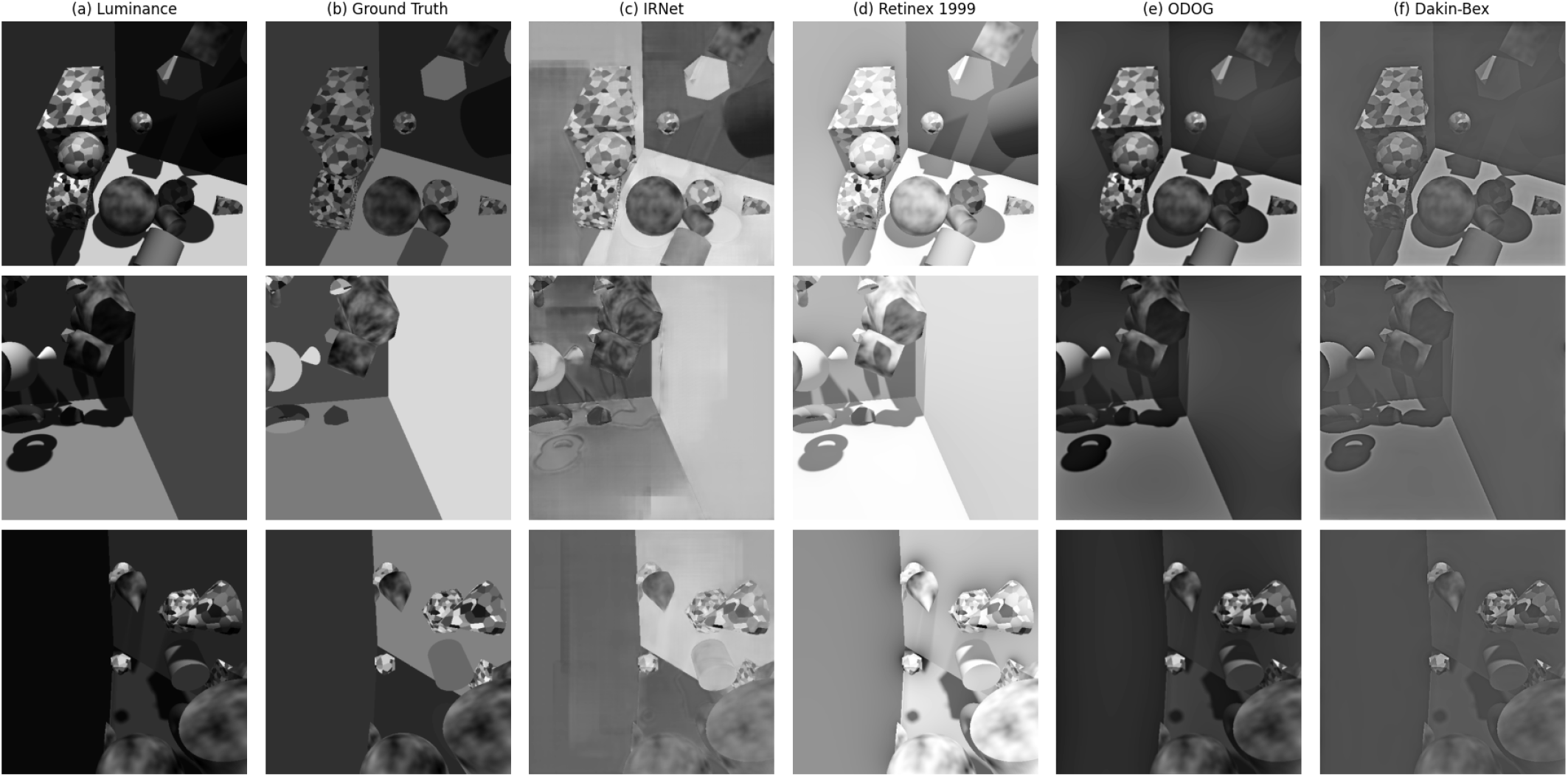
Typical images used for training and evaluating models, showing (a) luminance and (b) ground truth reflectance, as well as model outputs for (c) IRNet, (d) retinex, (e) ODOG, and (f) the Dakin-Bex model.

The network was trained via supervised learning. The inputs were the rendered scenes of geometric objects, and the target outputs were the reflectance maps. We used the ADAM optimizer (Kingma, 2014) with a mean-squared-error loss function. We used an initial learning rate of 0.001, and allowed ADAM to change it adaptively during training. Batch size was 64 images, and training continued for ten epochs. The loss typically plateaued between epochs 5 and 6. Training was carried out on two NVIDIA Tesla P100 GPUs.

#### Denoising control

To evaluate whether the network’s predictions of lightness illusions depended on the training objective, or were simply consequences of the network architecture, we trained the same network on a denoising task. We used the same training set of 100,000 grayscale images (Figure 5). We added white Gaussian noise with a pixelwise standard deviation of *σ* = 0.08 luminance units to the luminance images, corresponding to 10% of the median luminance across all images. We trained IRNet to output the corresponding noiseless luminance images, with a mean squared error (MSE) loss. To examine variability across training runs, we trained ten networks with different random seeds. Training ran for 10 epochs, with the average initial MSE of 0.021 decreasing to a final value of 0.002. The resulting illusion strengths are shown as dots in the dataset labeled ‘IRNet denoise’, with the corresponding bars showing means over the ten training runs, in Figure 6.

**Figure 6.**
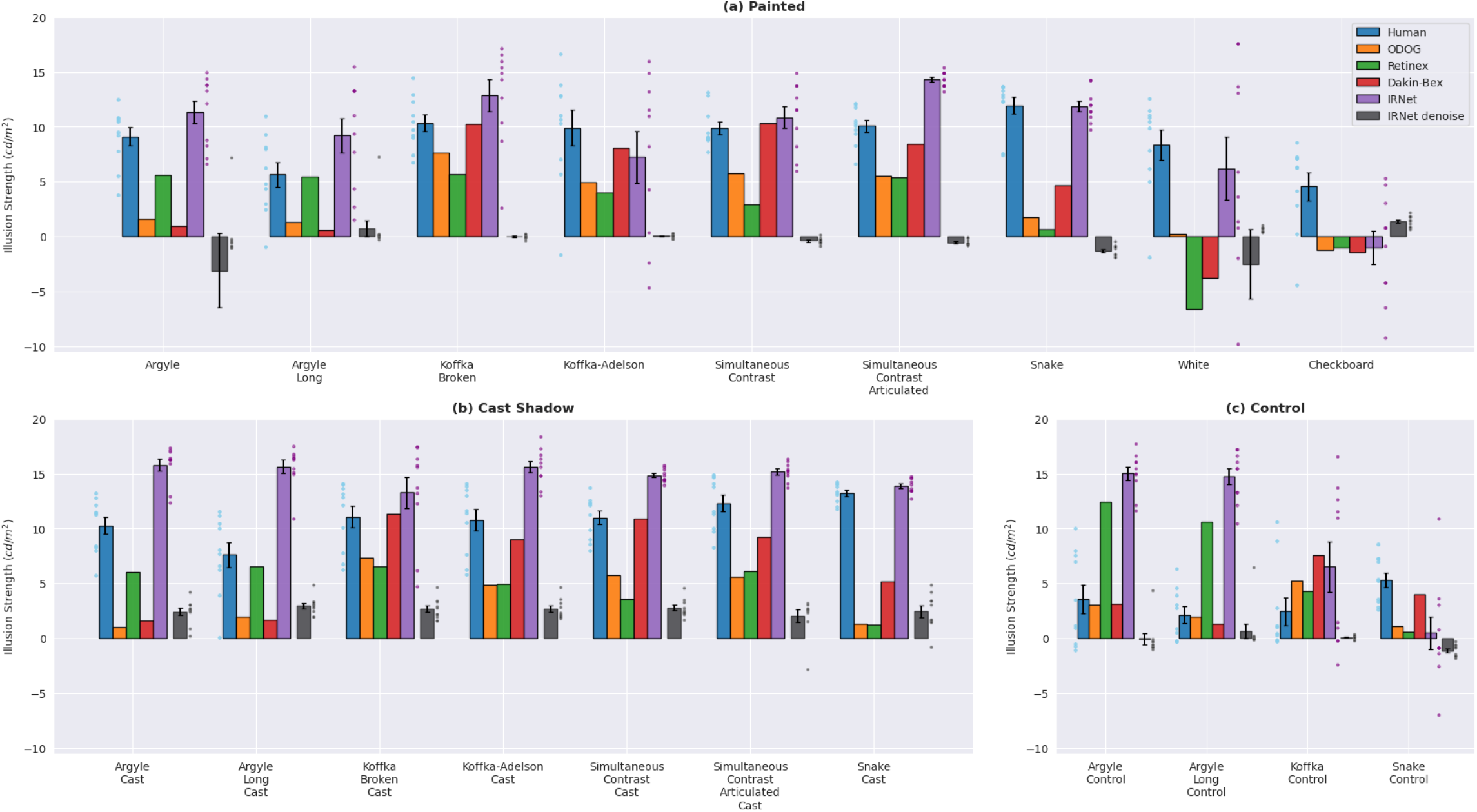
Illusion strength for human observers, IRNet, and classical models and IRNet trained for denoising, in (a) painted shadow images, (b) cast shadow images, and (c) control images. Dots next to the bars for humans and IRNet show results for individual observers and training runs, respectively. Error bars show standard errors, across observers for humans and across training runs for IRNet.

#### Classical models

To provide points of comparison for results with IRNet, we also tested three models of human lightness and brightness perception. Lightness is usually defined as perceived reflectance, and brightness as perceived luminance. However, the distinction between the two is often elided in the research literature, and given the small number of available computational models of lightness, we chose to include brightness models for comparison as well.

We tested the following models. Retinex is a lightness model that estimates reflectance by integrating a thresholded luminance gradient across an image (Land & McCann, 1971); we used one of McCann’s variants of this algorithm (Funt, Ciurea, & McCann, 2000; McCann, 1999). Dakin and Bex’s (2003) adaptive filtering algorithm models lightness and brightness by adjusting the Fourier amplitude spectrum of an image to match the 1*/f* ^*α*^ spectrum that is typical of natural images. ODOG is a brightness model that uses banks of elongated centre-surround receptive fields at a range of scales and orientations, followed by contrast normalization (Blakeslee & McCourt, 1999). As a baseline, we also tested a null model whose output was simply the luminance image provided as input, unchanged. None of these models give outputs in reflectance units, so we rescaled their outputs using an optimal affine transformation. We applied each model to the 10^5^ images in the training set, and found the affine transformation that gave each model the lowest median absolute error when predicting ground truth reflectance at 10^3^ pixels randomly sampled from each training image, for a total of 10^8^ pixels. In subsequent analyses, we applied these affine transforms to the output of each model, in order to convert the outputs in arbitrary units into reflectance estimates. To find these affine transformations, we used MATLAB’s ‘fminsearch’ minimization routine on an objective function that computed median absolute prediction error, with the slope and *y*-intercept of the affine transformation as free parameters.

We did not test the machine learning models reviewed in the Introduction that have been shown to reproduce lightness or brightness illusions, as none had code and trained model weights that were publicly available.

#### A simulated matching task

To examine the network’s and classical models’ responses to lightness illusions, we ran them in a simulation of the matching experiment reported above with human observers. We applied the network and models to the same stimuli shown to human observers (Figure 3), with a fixed reference patch reflectance and a range of match patch reflectances. For each image, we took the network’s and models’ match settings to be the match reflectance where the median pixelwise output over the match patch was the same as the median pixelwise output over the reference patch. These median outputs at the reference and match patches were never exactly the same for the finite number of test images used, so we found the match settings by interpolation or extrapolation. The match reflectances in the test images used were finely spaced, and only a small amount of extrapolation was required in a few cases, so interpolation and extrapolation were straightforward.

The Supporting Information includes the illusion images, trial-by-trial data for the human observers, an implementation of the deep learning network and training routine, code for classical lightness and brightness models, and code for the data analyses reported here (https://github.com/JaykishanPatel/irnet_supporting).

## Results

Figure 6 shows the results of the lightness matching task for human observers, IRNet, and the three classical models. In the following sections, we describe each group of results.

### Lightness illusions for human observers

The blue bars in Figure 6, on the left side of each group of bars, show the strength of each illusion, averaged over observers. The adjacent blue dots show the illusion strength for individual observers. Recall that we defined ‘illusion strength’ to be the reference luminance minus the match luminance, so positive values mean that observers chose a match luminance that was lower than the reference luminance. In each case, a positive value of illusion strength is what we would expect from previous literature. As expected, observers had a strong tendency to assign a lower luminance to the match region in the painted and cast-shadow stimuli, as indicated by the greater-than-zero illusion strength in these conditions (panels (a) and (b)). This tendency was much reduced for the control stimuli (panel (c)). Interestingly, there were large individual differences, with a few observers perceiving weak illusions or no illusions at all, even with these classic figures. Individual differences in lightness perception have been noted before (e.g., Murray, 2020; Patel et al., 2018), but have not been thoroughly explored (though see Brainard and Hurlbert (2015) and related work on colour constancy).

The first data column of Table 1 shows the results of one-tailed *t*-tests on illusion strength. We tested (a) whether the mean match luminance for each image was significantly lower than the reference luminance (the ‘match vs. reference’ rows in the table), and (b) whether the illusion was significantly stronger in the cast and painted shadow images than in the corresponding control image, for illusions that had control images (the ‘illusion vs. control’ rows). All comparisons were significant (*p <* 0.05, not corrected for multiple comparisons). These results show that, overall, human observers perceived the expected illusions in these novel versions of classic figures, and perceived stronger illusions than in the control conditions. Note that Table 1 reports results for each painted and cast shadow condition, and also for some comparisons between those conditions and control conditions. As a result, there is not a one-to-one correspondence between rows in Table 1 and conditions in Figure 6.

**Table 1.**
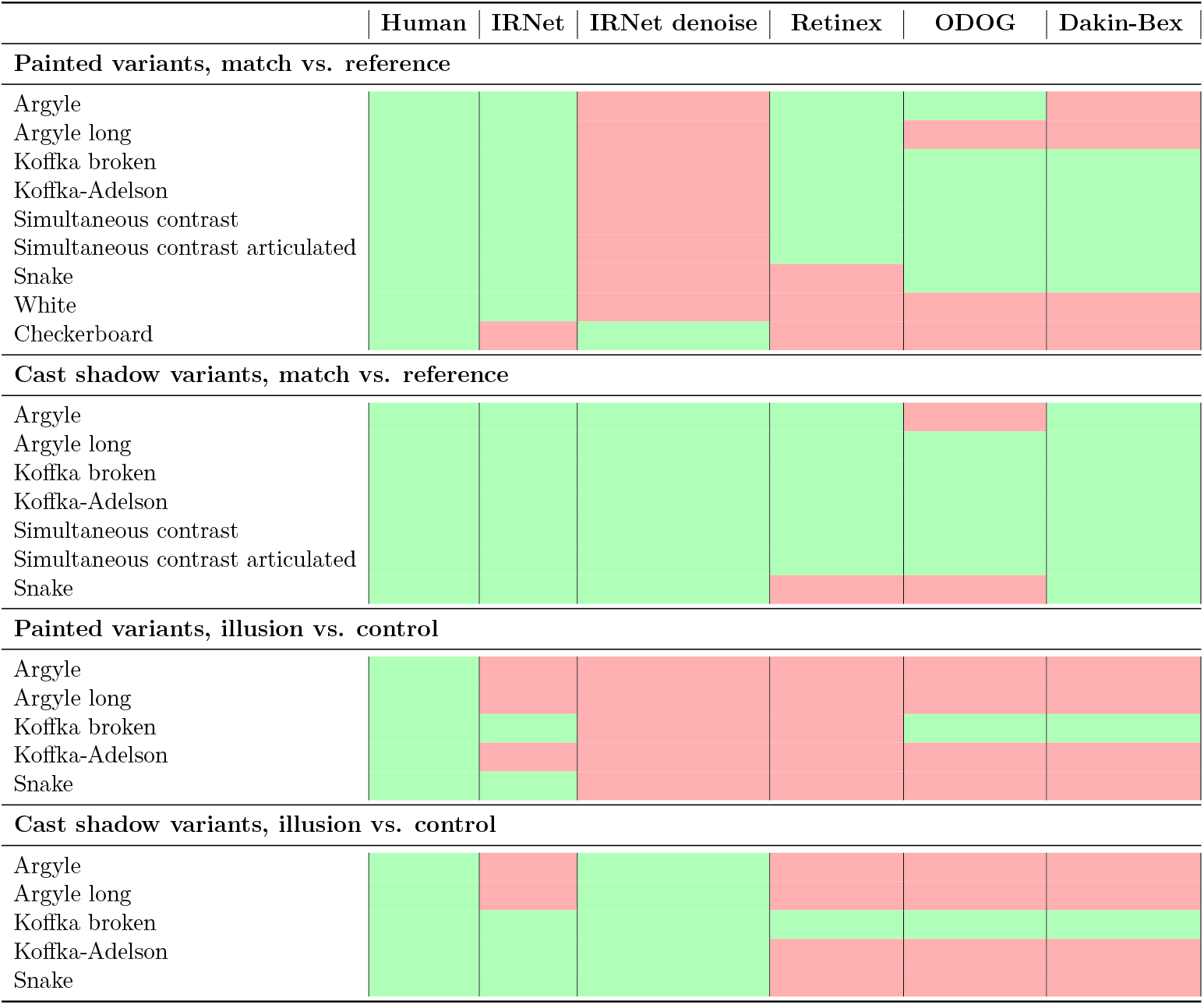
Tests of illusion effects for human and model observers. For human observers, IRNet and denoising IRNet (first three columns), green cells indicate a significant effect (p < 0.05) and red cells indicate no significant effect. For the illusion vs. control comparisons, we used paired t-tests. For the classical models (remaining three columns), green cells indicate an illusion effect of at least 5%, and red cells indicate no such effect. See the main text for details.

### Overview of network performance

Before reporting IRNet’s results with the illusion figures, we give a brief overview of its performance at reflectance estimation after training. The untrained model (random initialization) produced an average mean squared error (MSE) of 0.45, whereas the model trained for 10 epochs reached an average MSE of 0.08. This reflects substantial learning. The error dropped markedly within the first 5–6 epochs and then plateaued. These error values were computed over 5,000 held-out test images.

Figure 5(a) shows typical test images, which were generated in the same manner as the training images, but were not used during training. Figure 5(b) shows the corresponding ground truth reflectance images. Figure 5(c) shows the network’s output in response to the test images in column (a). The network was clearly able to remove or reduce lighting-related features such as shadows and shading, both on the geometric objects and on the walls. There was some residual mottling, which is particularly visible in regions of uniform reflectance.

For comparison, the remaining columns of Figure 5 show results from the three classical models. These models did not remove the effects of illumination to as large an extent as the network, with shadows and shading remaining clearly visible in their outputs.

Figure 7(a) shows a scatterplot of the network’s output versus ground truth reflectance, for test images not used during training. There was a strong positive correlation (*r* = 0.81), averaged over 10 training runs, between the model’s output and true reflectance. The average median absolute reflectance error over the 10 runs was 0.08, on a reflectance scale that ranges from 0 to 1. These values are based on the reflectance outputs for each training run from 10^3^ randomly selected pixels from each of the 5 *×* 10^3^ test images, for a total of 5 *×* 10^6^ pixels. The scatterplot shows 10^3^ pixels randomly sampled from this larger set.

**Figure 7.**
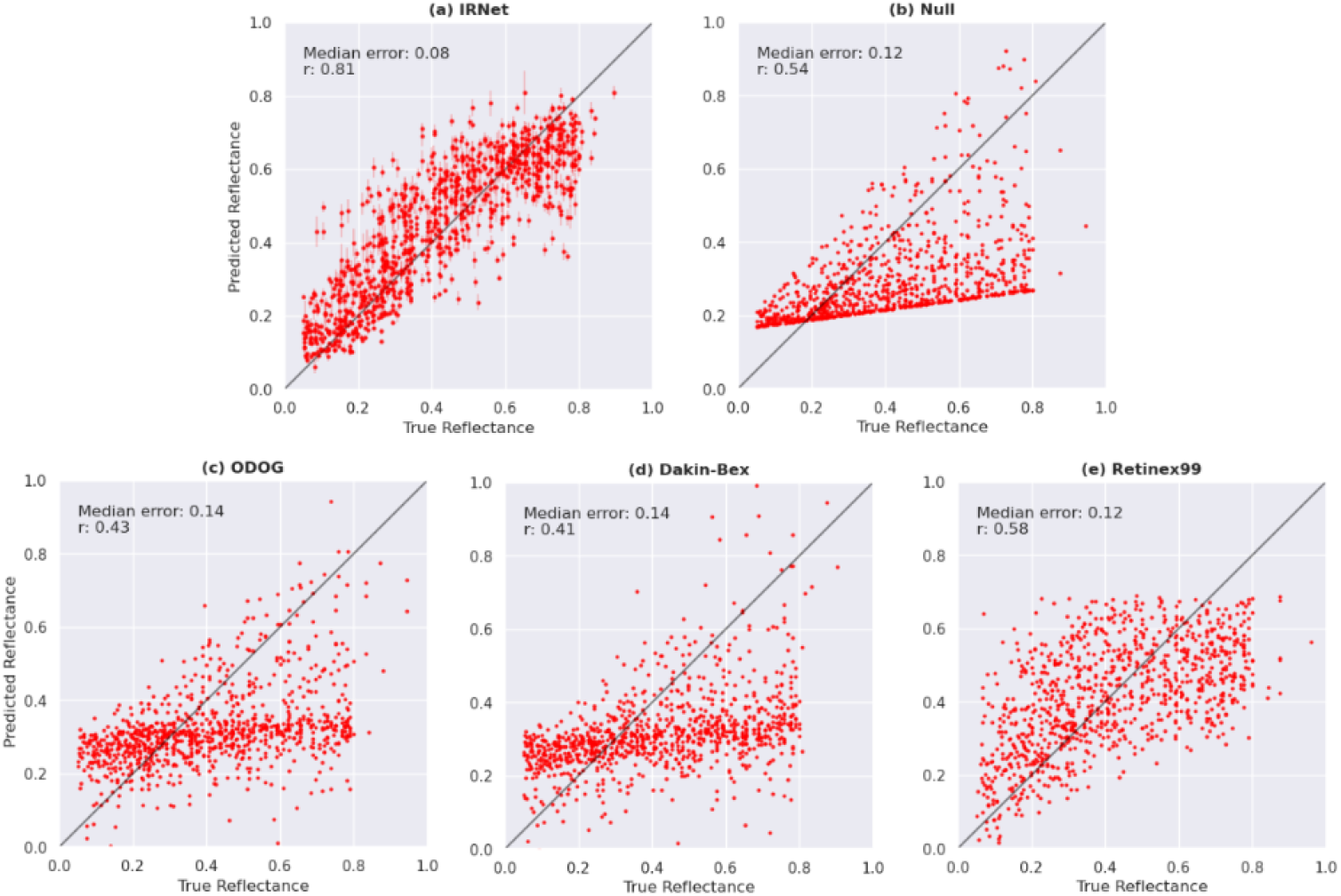
Scatterplots of the models’ reflectance predictions against ground truth reflectance. Error bars in (a) show the standard deviation across training runs of IRNet.

Figure 7 also shows scatterplots for the classical models, as well as the null model whose output was simply an optimized affine transform of the luminance input image. The correlation of these models’ outputs with true reflectance was much lower than for IRNet, and the median absolute reflectance error was higher. This error metric reflects model performance on test data, based on pixelwise predictions across 5 million randomly sampled test points. Note that this pixelwise median absolute error differs from the mean squared error used as the training loss. These classical models tended to overestimate low reflectances and underestimate high reflectances. The most successful classical model was retinex, which had a median reflectance error of 0.12, the same as the null model. The ODOG and Dakin-Bex models were outperformed even by the null model. As noted above, not all of these models were intended by their creators to be models of lightness. Retinex is a lightness model, but ODOG is a brightness model, and the Dakin-Bex model is sometimes described as a lightness model and sometimes as a brightness model.

### Lightness illusions for IRNet

The purple bars in Figure 6, the second from the right side in each group, show the average illusion strength for IRNet across training runs. The adjacent dots show the illusion strength for individual training runs. The network manifests illusions that are broadly consistent with the human matching data reported above, though with some differences. We first describe results for individual classes of illusions, and then report statistical tests similar to those we reported for human observers.

The network shows strong illusions in the argyle and long-range argyle illusions, and they are somewhat stronger in the cast shadow conditions than in the painted shadow conditions, which is consistent with human matching data. However, the network’s illusion is about equally strong in these conditions and in the argyle control conditions, which is not consistent with human data. Previous studies have also found the argyle illusion to be particularly challenging to model (Blakeslee & McCourt, 2012; Murray, 2020).

For the Koffka group of illusions, the network’s outputs are largely consistent with human matching data. The network manifests strong illusions in the painted and cast shadow variants of the Koffka-broken illusion. For the Koffka-Adelson illusion, it shows a strong effect in the cast shadow variant, consistent with human matches, and shows a weaker, but still significant, illusion in the painted shadow variant. This weakening in the painted shadow condition is not consistent with human matches. The network shows a weaker illusion in the Koffka control figure, which is directionally consistent with human matches. The illusion in the control condition is, however, stronger for the network than for human observers.

The network shows strong illusions in the cast and painted shadow variants of the simultaneous contrast and articulated simultaneous contrast figures. It also shows strong illusions in the painted and cast shadow variants of the snake illusion, and a much weaker effect in the control condition. These effects are all consistent with human matches.

The network’s illusion strength is negative for the checkerboard figure, which is not consistent with human matching data. The network’s illusion is in the correct direction (and significantly greater than zero) for White’s illusion, but highly variable across training runs. These are the two *assimilation* illusions in this stimulus set, meaning that for human observers, the reference and match patches tend to appear more similar to their surrounds, rather than less similar as in contrast illusions.

To summarize these findings and test for significant effects, the second data column of Table 1 shows the results of the same significance tests we reported for human observers, evaluated across training runs. These tests show that the network exhibited many of the same illusion effects as human observers, the exceptions being the checkerboard illusion, and the illusion vs. control comparison for the argyle and Koffka-Adelson illusions. The network produced illusion effects even in the painted images, despite being trained for reflectance estimation. We did not correct for multiple comparisons, so there may be a small number of type I errors, but overall the qualitative similarity between results for human observers and IRNet is clear.

### Lightness illusions for classical models

Figure 6 also shows illusion strength for the classical models, in the middle three bars of each group. These models predicted some illusions in the painted and cast shadow conditions, though not all, and a particular failure mode was that they often predicted even stronger illusions in the control conditions. This is consistent with Murray’s (2020) findings with these models and simpler 2D versions of the illusions.

The three rightmost columns of Table 1 summarize the illusions found in the classical models’ responses. The models had deterministic outputs, so significance tests were not appropriate. Instead, we considered an illusion to be present in the ‘match vs. reference’ tests if the match luminance was at least 5% less than the reference reflectance, and in the ‘illusion vs. control’ tests if the match luminance was at least 5% lower in the illusion stimulus than in the control stimulus. These low, near-threshold criteria give a generous interpretation of whether an illusion was present in each case.

Even with these liberal criteria, the models failed to account for several lightness illusions, and as noted above, usually predicted an equally strong or stronger illusion in the control conditions.

## Discussion

Lightness perception is a long-standing area of research, and many studies have revealed important principles of lightness perception (Adelson, 2000; Arend & Spehar, 1993; Brainard & Maloney, 2011; Gilchrist, 2006; Murray, 2021). However, this work has resulted in few general-purpose, image-computable models. Here we found that a convolutional neural network, using a common architecture and trained for reflectance estimation, is a promising starting point for such a model.

One of our goals was to evaluate a specific CNN as a model of lightness perception. We found that IRNet has several strengths as a model. It is image-computable, and can be applied to arbitrary luminance images. Its outputs are absolute reflectance values that do not require normalization. In our tests, it discounted lighting-related effects such as shadows and shading, and its reflectance estimates were strongly correlated with ground truth. It had some ability to generalize beyond its training set, producing reasonable outputs on the illusion images, which were clearly not samples from the training set. It assigned different reflectances to isoluminant test regions in most of the illusion images we tested, and these differences were smaller with most of the control stimuli. The matching behavior of the network largely followed the behaviour of human observers in a corresponding matching experiment. Furthermore, the network exhibited these illusions even though it was not trained to do so; it was simply trained to estimate reflectance as well as possible from luminance images. On these measures, the network far outperformed the three classical models we tested.

The primary focus of this work was to evaluate the behavioral similarity between the network and human observers. While the network does not provide direct insight into the computations underlying lightness perception, its outputs align with human judgments across a range of conditions and outperform those of classical models. A key limitation of using deep learning architectures as models of the human visual system is their black-box nature. Although this work does not reveal how the brain computes lightness, it demonstrates that such models can exhibit strikingly human-like patterns of behavior when trained on appropriate tasks. This finding suggests that certain aspects of lightness perception, such as illusions, may emerge as a by-product of creating a perceptual correlate of reflectance. Future work may investigate the extent to which unsupervised objectives yield similar behaviors (e.g., Storrs, Anderson, and Fleming (2021)), or the internal structure of the network (for example through saliency analysis or lesion-based ablation studies). Here we take a functional perspective, emphasizing the value of image-computable models that replicate human-like behavior in controlled perceptual tasks.

The network also has weaknesses as a model. It did not account for two assimilation illusions. This was not entirely surprising, as assimilation illusions are difficult to explain in terms of discounting illumination. The snake illusion, for example, is a contrast illusion that contains isoluminant diamonds in bright and dark surrounds (Figure 1). If an observer believes that the surrounds have different illumination levels, a logical conclusion is that the diamonds in the brighter, more strongly illuminated region must have a lower reflectance. Thus illumination discounting offers a straightforward explanation of contrast-like effects. It is less obvious, though, how illumination discounting can make a test patch appear *more* similar to its surround, as in assimilation illusions.

The network also failed to account for the argyle illusion, predicting a stronger illusion in the control condition. The reason for this is unclear, but it may be that the argyle figure was too dissimilar to the reflectance patterns in the training set, e.g., images in the training set did not have the thin lines that cross between light and dark regions in the argyle figure. However, as noted earlier, it is also true that the argyle illusion has confounded previous models that have been able to account for a range of other illusions (Blakeslee & McCourt, 2012; Murray, 2020).

Figure 6 also shows that, although the network usually predicted the correct direction of illusory effects for human observers, the effects were often much stronger for the network than for humans, particularly in the cast shadow conditions. This is supported by Figure 8, which scatterplots the illusion strength for IRNet and humans. In 15 of the 20 conditions, illusions were stronger for IRNet, which is significantly greater than chance according to a binomial test (*p* < 0.01).

**Figure 8.**
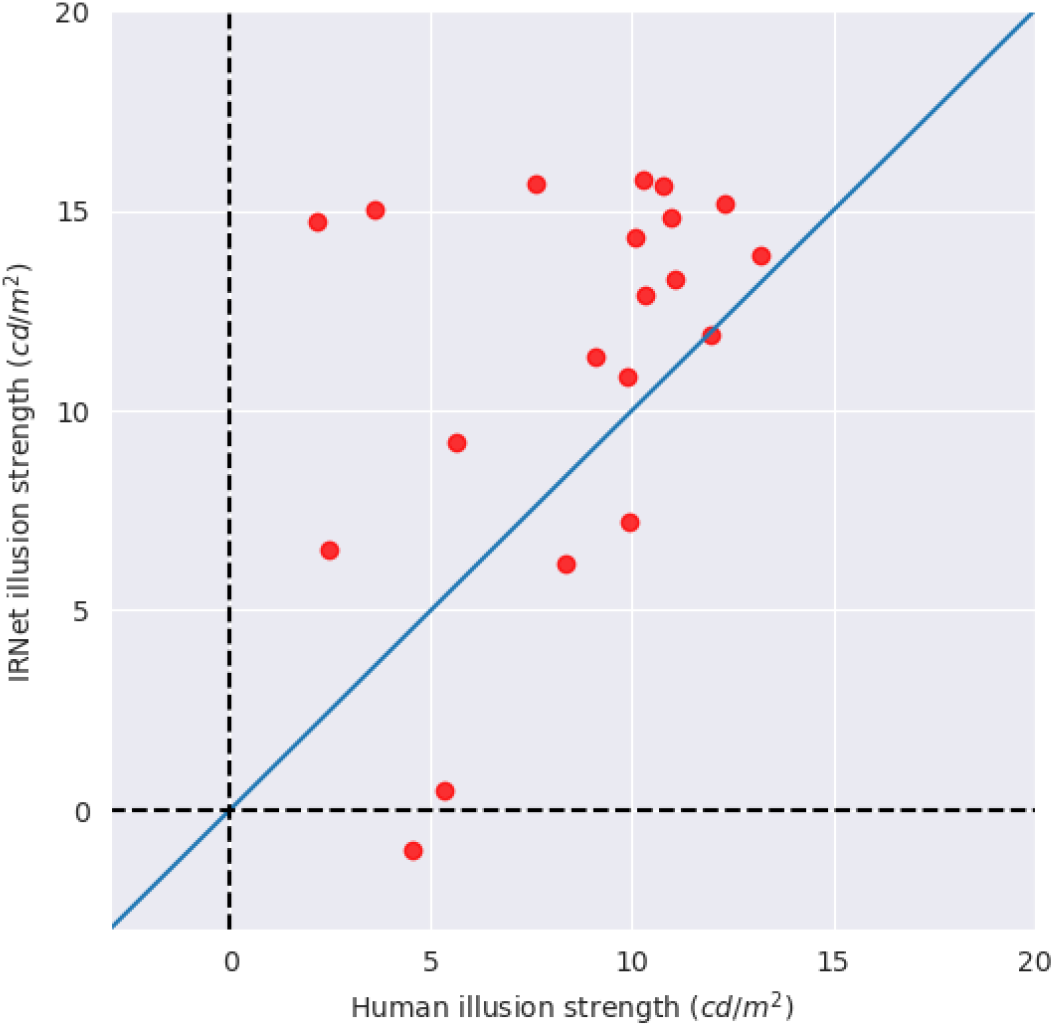
Illusion strength for human observers and IRNet. The diagonal blue line shows *y* = *x*. Illusions tended to be stronger for the network than for human observers.

We have interpreted our findings as showing that lightness illusions can be a byproduct of training for reflectance estimation. However, previous work has shown that the choice of network architecture can strongly bias the behaviour of trained networks (Ulyanov, Vedaldi, & Lempitsky, 2018). In our case, might it be that the IRNet architecture itself is largely responsible for our results, and that strong lightness illusions would emerge even when the network is trained on other tasks? To address this possibility, we trained the IRNet architecture on a denoising task, using the same training images we used for reflectance estimation, and tested the trained network for lightness illusions. Figure 6 shows that any resulting lightness illusions were very weak, which supports the notion that training on reflectance estimation played an essential role in our findings.

We did not systematically evaluate the network’s ability to generalize beyond its training set, but specialized training methods are often needed to produce deep learning networks that generalize well (Srivastava, Hinton, Krizhevsky, Sutskever, & Salakhutdinov, 2014; Zhang, Bengio, Hardt, Recht, & Vinyals, 2017). Although the illusion images were not samples from the training set, they were nevertheless similar in many ways. We do not expect that the network would perform well on more realistic, natural images, and this is another way in which the network will need further development.

Perhaps the most important limitation of the network in its current form is that it provides little insight into how lightness perception is achieved algorithmically, either by human observers or by the network itself. Our results are more informative about the relationship between natural scene statistics and lightness perception, than about the computations that underlie lightness perception. In effect, we treat the network as a trainable inference engine, and observe what behaviours emerge when it is trained in a controlled virtual environment that shares some features with the real world. This is a productive approach to understanding vision, but in future work it would also be useful to explore methods for understanding the features exploited by convolutional networks, or to investigate alternative architectures that are more easily interpreted (Dosovitskiy, 2020; Selvaraju et al., 2020; Simonyan, 2013; Zeiler, 2014; Zhou, Khosla, Lapedriza, Oliva, & Torralba, 2016). Another promising direction for future work is to examine whether alternative architectures, such as shallower CNNs, recurrent neural networks, or transformer-based models, trained on the same reflectance estimation task and dataset, produce similar or different patterns of behavior. To support such work, we have made our code for training and analysis publicly available (https://github.com/JaykishanPatel/irnet_supporting).

Another of our goals was to evaluate the notion that many phenomena in lightness perception, including some strong lightness illusions, are byproducts of the visual system’s attempt to compute a stable correlate of surface reflectance from ambiguous 2D images. The network’s successes suggest that this is a useful approach to understanding lightness. The training set consisted of achromatic renderings of a few kinds of simple geometric objects and surface patterns, and yet a network trained on this dataset was susceptible to several of the same strong lightness illusions as human vision. Preliminary work suggests that even in non-illusory stimuli, such networks also have partial failures of lightness constancy that are similar to those of human observers (Flachot et al., 2025).

All papers reviewed in the ‘Previous work’ section above (Gomez-Villa et al., 2020; Kubota et al., 2021; Mukherjee et al., 2024), with the exception of Corney and Lotto (2007), examined networks trained for denoising, deblurring, or both. From those studies we might conclude that lightness illusions are byproducts of low-level image enhancement processes. That may be partly the case, but our findings show that even without such low-level processes, strong illusions can emerge simply from an attempt to infer reflectance from luminance images. One interesting approach to examining connections between previous work and the present work would be to incorporate known features of early visual processing, such as optical blur and varying photoreceptor density, or other constraints on visual processing (Cottaris, Wandell, Rieke, & Brainard, 2020), into a normative model of reflectance estimation such as ours.

Understanding lightness illusions as a byproduct of rational processes for estimating albedo suggests that the question of what constitutes an ‘illusion’ is more nuanced than it might appear at first. For example, in cast shadow version of the Koffka-Adelson figure (Figure 3(b)), the semicircle on the darker side of the shadow appears lighter than the isoluminant semicircle on the other side. Another way of describing the same percept, however, is that the higher-albedo semicircle depicted on the shadowed side of the boundary appears lighter than the darker-albedo semicircle depicted on the other side. Clearly, then, this figure is only an illusion if we expect the relevant percept to be correlated with image luminance. If we expect it to be correlated with albedo, then far from being an illusion, this figure demonstrates the human visual system’s remarkable ability to use ambiguous information to correctly estimate the albedo of a depicted surface.

Interestingly, it may be that only networks in some intermediate range of power are good models of lightness perception. In the ‘painted’ illusion figures we used, the illusions seem to depend on the observer misinterpreting a light-dark reflectance boundary as a shadow boundary. The network we tested here apparently did so as well, and assigned similar reflectances to the painted and cast shadow variants of illusions.

However, there are cues in the painted figures that would presumably allow a sufficiently thorough observer to correctly interpret the boundary as a reflectance edge, e.g., a light-dark boundary on one side of the cube does not continue onto an adjacent side, as a shadow boundary usually would. A more powerful or more thoroughly trained network might not be susceptible to illusions in the painted figures, and so would be a weaker model of human vision.

Indeed, it is noteworthy that the network is subject to lightness illusions in the painted conditions at all. One might have thought that in order for a network to be reasonably good at estimating surface reflectance, it would have to distinguish between paint and shadow well enough to not be deceived by the painted figures. Such a finding would undercut an explanation of these illusions as a natural byproduct of estimating reflectance. However, this was not the case, and so this explanation remains viable.

## Acknowledgements

This research was co-funded by the following sources. (a) NSERC Canada Graduate Scholarship - Doctoral to JP. (b) A postdoctoral fellowship to AF from the Canada First Research Excellence Fund, through the Vision: Science to Applications program at York University. (c) European Union funding to TSAW (ERC, SEGMENT, 101086774). Views and opinions expressed are however those of the author(s) only and do not necessarily reflect those of the European Union or the European Research Council. Neither the European Union nor the granting authority can be held responsible for them. This work was supported by the Deutsche Forschungsgemeinschaft (German Research Foundation, DFG) under Germany’s Excellence Strategy (EXC 3066/1 “The Adaptive Mind”, Project No. 533717223). (d) An NSERC Discovery Grant to RFM. (e) Computing resources were provided by SHARCNET (www.sharcnet.ca) and the Digital Research Alliance of Canada (alliancecan.ca).

## References

Adelson, E. H. (1993). Perceptual organization and the judgment of brightness. Science, 262 (5142), 2042–2044.

Adelson, E. H. (2000). Lightness perception and lightness illusions. In M. Gazzaniga (Ed.), The new cognitive neurosciences (pp. 339–351). Cambridge, MA: The MIT Press.

Allred, S. R., & Brainard, D. H. (2013). A Bayesian model of lightness perception that incorporates spatial variation in the illumination. Journal of Vision, 13(7):18, 1–18.

Arend, L. E., & Spehar, B. (1993). Lightness, brightness, and brightness contrast: 1. illuminance variation. Perception & Psychophysics, 54 (4), 446–456.

Barron, J. T., & Malik, J. (2015). Shape, illumination, and reflectance from shading. IEEE Transactions on Pattern Analysis and Machine Intelligence, 37 (8), 1670–1687.

Barrow, H., Tenenbaum, J., Hanson, A., & Riseman, E. (1978). Recovering intrinsic scene characteristics. Computer Vision Systems, 2 (3-26), 2.

Belhumeur, P. N., Kriegman, D. J., & Yuille, A. L. (1999). The bas-relief ambiguity. International Journal of Computer Vision, 35 (1), 33–44.

Blakeslee, B., & McCourt, M. E. (1999). A multiscale spatial filtering account of the white effect, simultaneous brightness contrast and grating induction. Vision Research, 39 (26), 4361–4377.

Blakeslee, B., & McCourt, M. E. (2012). When is spatial filtering enough? Investigation of brightness and lightness perception in stimuli containing a visible illumination component. Vision Research, 60, 40–50.

Blender 2.92 Documentation Team. (2021). Blender [Computer software manual]. Blender Foundation, Amsterdam. Retrieved from https://docs.blender.org/manual/en/2.92

Brainard, D. H., & Freeman, W. T. (1997). Bayesian color constancy. JOSA A, 14 (7), 1393–1411.

Brainard, D. H., & Hurlbert, A. C. (2015). Colour vision: understanding# thedress. Current Biology, 25 (13), R551–R554.

Brainard, D. H., Longere, P., Delahunt, P. B., Freeman, W. T., Kraft, J. M., & Xiao, B. (2006). Bayesian model of human color constancy. Journal of Vision, 6(11):10. doi: 10.1167/6.11.10

Brainard, D. H., & Maloney, L. T. (2011). Surface color perception and equivalent illumination models. Journal of Vision, 11(5):1. doi: 10.1167/11.5.1

Corney, D., & Lotto, R. B. (2007). What are lightness illusions and why do we see them? PLoS Computational Biology, 3 (9), e180.

Cottaris, N. P., Wandell, B. A., Rieke, F., & Brainard, D. H. (2020). A computational observer model of spatial contrast sensitivity: effects of photocurrent encoding, fixational eye movements, and inference engine. Journal of Vision, 20(7):17, 1-25.

Dakin, S. C., & Bex, P. J. (2003). Natural image statistics mediate brightness ‘filling in’. Proceedings of the Royal Society of London. Series B: Biological Sciences, 270 (1531), 2341–2348.

DeValois, R. L., & DeValois, K. K. (1988). Spatial vision. Oxford, UK: Oxford University Press.

Dosovitskiy, A. (2020). An image is worth 16×16 words: Transformers for image recognition at scale. arXiv preprint arXiv:2010.11929.

Flachot, A., Patel, J., Wallis, T. S. A., Brubaker, M. A., Brainard, D. H., & Murray, R. F. (2025). Deep neural networks trained for estimating albedo and illumination achieve lightness constancy differently than human observers. bioRxiv doi: 10.1101/2025.07.10.664065.

Funt, B., Ciurea, F., & McCann, J. J. (2000). Retinex in MATLAB. In Proceedings of the IS&T/SID Eighth Color Imaging Conference (p. 112–121). Scottsdale, AZ, November 2000.

Gilchrist, A. (2006). Seeing black and white. New York: Oxford University Press.

Gomez-Villa, A., Martín, A., Vazquez-Corral, J., Bertalmío, M., & Malo, J. (2020). Color illusions also deceive cnns for low-level vision tasks: Analysis and implications. Vision Research, 176, 156–174.

Gomez-Villa, A., Martín, A., Vazquez-Corral, J., Bertalmío, M., & Malo, J. (2022). On the synthesis of visual illusions using deep generative models. Journal of Vision, 22 (8), 2, 1–18.

Hess, C., & Pretori, H. (1894/1970). Quantitative investigation of the lawfulness of simultaneous brightness contrast. Perceptual and Motor Skills, 31 (Suppl. 2-V31), 947–969.

Katz, D. (1935). The world of colour. London: Kegan & Paul.

Kingdom, F. A. A. (2008). Perceiving light versus material. Vision Research, 48 (20), 2090–2105. doi: 10.1016/j.visres.2008.03.020

Kingdom, F. A. A. (2011). Lightness, brightness and transparency: a quarter century of new ideas, captivating demonstrations and unrelenting controversy. Vision Research, 51, 652–673.

Kingma, D. P. (2014). Adam: A method for stochastic optimization. arXiv preprint arXiv:1412.6980.

Koffka, K. (1935). Principles of gestalt psychology. New York: Harcourt, Brace, & World, Inc.

Kubota, Y., Hiyama, A., & Inami, M. (2021). A machine learning model perceiving brightness optical illusions: quantitative evaluation with psychophysical data. In Proceedings of the Augmented Humans International Conference (p. 174–182). Association for Computing Machinery.

Land, E. H., & McCann, J. J. (1971). Lightness and retinex theory. Journal of the Optical Society of America, 61 (1), 1–11.

Li, Z., & Chen, J. (2015, June). Superpixel segmentation using linear spectral clustering. In Proceedings of the IEEE Conference on Computer Vision and Pattern Recognition.

Li, Z., Yu, T.-W., Sang, S., Wang, S., Song, M., Liu, Y., … others (2021). Openrooms: An open framework for photorealistic indoor scene datasets. In Proceedings of the IEEE/CVF Conference on Computer Vision and Pattern Recognition (pp. 7190–7199).

McCann, J. (1999). Lessons learned from mondrians applied to real images and color gamuts. In Color and Imaging Conference (Vol. 1999, pp. 1–8).

McCluney, R. (1994). Introduction to radiometry and photometry. Artech House, Inc.

Mukherjee, A., Paul, A., & Ghosh, K. (2024). Deep learning models for perception of brightness related illusions. Applied Intelligence, 54, 10259–10283.

Murray, R. F. (2013). Human lightness perception is guided by simple assumptions about reflectance and lighting. In B. E. Rogowitz, T. N. Pappas, & H. de Ridder (Eds.), Proceedings of SPIE 8651, Human Vision and Electronic Imaging XVIII.

Murray, R. F. (2020). A model of lightness perception guided by probabilistic assumptions about lighting and reflectance. Journal of Vision, 20(7):28. doi: 10.1167/jov.20.7.28

Murray, R. F. (2021). Lightness perception in complex scenes. Annual Review of Vision Science, 7, 417–436.

Paszke, A., Gross, S., Massa, F., Lerer, A., Bradbury, J., Chanan, G., … others (2019). Pytorch: An imperative style, high-performance deep learning library. Advances in Neural Information Processing Systems, 32.

Patel, K., Munasinghe, A., & Murray, R. (2018). Lightness matching and perceptual similarity. Journal of Vision, 18 (5), 1–1.

Peirce, J., Gray, J. R., Simpson, S., MacAskill, M., Höchenberger, R., Sogo, H., … Lindeløv, J. K. (2019). Psychopy2: Experiments in behavior made easy. Behavior Research Methods, 51, 195–203.

Purves, D., & Lotto, R. B. (Eds.). (2010). Why we see what we do redux: a wholly empirical theory of vision. Sinauer Associates.

Rother, C., Kiefel, M., Zhang, L., Schölkopf, B., & Gehler, P. (2011). Recovering intrinsic images with a global sparsity prior on reflectance. Advances in Neural Information Processing Systems, 24.

Sato, S., Kaneko, T., Murasaki, K., Yoshida, T., Tanida, R., & Kimura, A. (2024). Unsupervised intrinsic image decomposition with lidar intensity enhanced training. arXiv preprint arXiv:2403.14089.

Schirillo, J. A. (2013). We infer light in space. Psychonomic Bulletin & Review, 20 (5), 905–915. doi: 10.3758/s13423-013-0408-1

Selvaraju, R. R., Cogswell, M., Das, A., Vedantam, R., Parikh, D., & Batra, D. (2020). Grad-cam: visual explanations from deep networks via gradient-based localization. International Journal of Computer Vision, 128, 336–359.

Shen, L., Yeo, C., & Hua, B.-S. (2013). Intrinsic image decomposition using a sparse representation of reflectance. IEEE Transactions on Pattern Analysis and Machine Intelligence, 35 (12), 2904–2915.

Simonyan, K. (2013). Deep inside convolutional networks: Visualising image classification models and saliency maps. arXiv preprint arXiv:1312.6034.

Srivastava, N., Hinton, G., Krizhevsky, A., Sutskever, I., & Salakhutdinov, R. (2014). Dropout: a simple way to prevent neural networks from overfitting. The journal of machine learning research, 15 (1), 1929–1958.

Storrs, K. R., Anderson, B. L., & Fleming, R. W. (2021). Unsupervised learning predicts human perception and misperception of gloss. Nature Human Behaviour, 5 (10), 1402–1417.

The MathWorks Inc. (2022). Matlab version: 9.13.0 (r2022b). Natick, Massachusetts, United States: The MathWorks Inc. Retrieved from https://www.mathworks.com

Ulyanov, D., Vedaldi, A., & Lempitsky, V. (2018). Deep image prior. In Proceedings of the IEEE Conference on Computer Vision and Pattern Recognition (CVPR) 2018.

Weiss, Y. (2001). Deriving intrinsic images from image sequences. In Proceedings Eighth IEEE International Conference on Computer Vision. ICCV 2001 (Vol. 2, p. 68–75 vol.2).

White, M. (1979). A new effect of pattern on perceived lightness. Perception, 8 (4), 413–416.

Yu, B., Yang, S., Cui, X., Dong, S., Chen, B., & Shi, B. (2023). Milo: Multi-bounce inverse rendering for indoor scene with light-emitting objects. IEEE Transactions on Pattern Analysis and Machine Intelligence, 45 (8), 10129–10142.

Yu, Y., & Smith, W. A. P. (2018). Inverserendernet: Learning single image inverse rendering. CoRR, abs/1811.12328.

Zeiler, M. (2014). Visualizing and understanding convolutional networks. In European conference on computer vision (Vol. 1311).

Zhang, C., Bengio, S., Hardt, M., Recht, B., & Vinyals, O. (2017). Understanding deep learning requires rethinking generalization.

Zhou, B., Khosla, A., Lapedriza, A., Oliva, A., & Torralba, A. (2016). Learning deep features for discriminative localization. In Proceedings of the ieee conference on computer vision and pattern recognition (pp. 2921–2929).

